# Isolation, extraction, and physicochemical characterization of exosome-like nanovesicles from ginseng adventitious roots in tissue culture

**DOI:** 10.64898/2026.10.12.758673

**Authors:** Leyi Xiao, Yuanzhu Liu, Chi Xiong, Cheng Sun, Junquan Cheng, Junxia Xiao, Yuping Rong, Rongchun Han, Qinying Yang, Xiao Liu, Rui Li, Cuiling Zhang, Xin Wan, Xianqian Liu, Jihong Huang

## Abstract

This study aimed to establish a method for isolating exosome-like nanovesicles from ginseng adventitious roots and their culture supernatant, to investigate the effects of methyl jasmonate (MeJA) induction on vesicle yield and composition, and to provide experimental evidence for the scalable production of ginseng-derived exosome-like nanovesicles. Exosome-like nanovesicles were isolated from ginseng adventitious roots and culture supernatant using low-speed centrifugation combined with stirred-cell ultrafiltration. Physicochemical characterization was performed using transmission electron microscopy (TEM), nanoparticle tracking analysis (NTA), Zeta potential analysis, and Q-ToF LC-MS lipidomics. Ginseng adventitious roots were treated with 200 μM MeJA to evaluate changes in vesicle particle concentration, total protein, total saponins, and miRNA expression before and after induction. Paraffin section fluorescence staining was used to observe the distribution of vesicle-related signals in adventitious root tissues. Exosome-like nanovesicles with typical bilayer membrane structures were isolated from both ginseng adventitious roots and culture supernatant.The particle size ranged from 77.333 to 171.733 nm, and the particle concentration ranged from 2.1E+8 to 7.6E+10 /mL. The vesicles contained lipid components including glycerides, phospholipids, and sphingolipids. Induction with 200 μM MeJA increased vesicle particle concentration and total protein content, significantly upregulated the total saponin level in vesicles derived from adventitious roots, and altered the miRNA expression pattern of the vesicles. Tissue section fluorescence observation showed overlap between vacuolar markers and membrane vesicle dye signals. Low-speed centrifugation combined with ultrafiltration can effectively prepare ginseng-derived exosome-like nanovesicles. MeJA induction can regulate vesicle yield and active components, providing foundational data for the scalable process development of such vesicles.

**Graphical Abstract:** 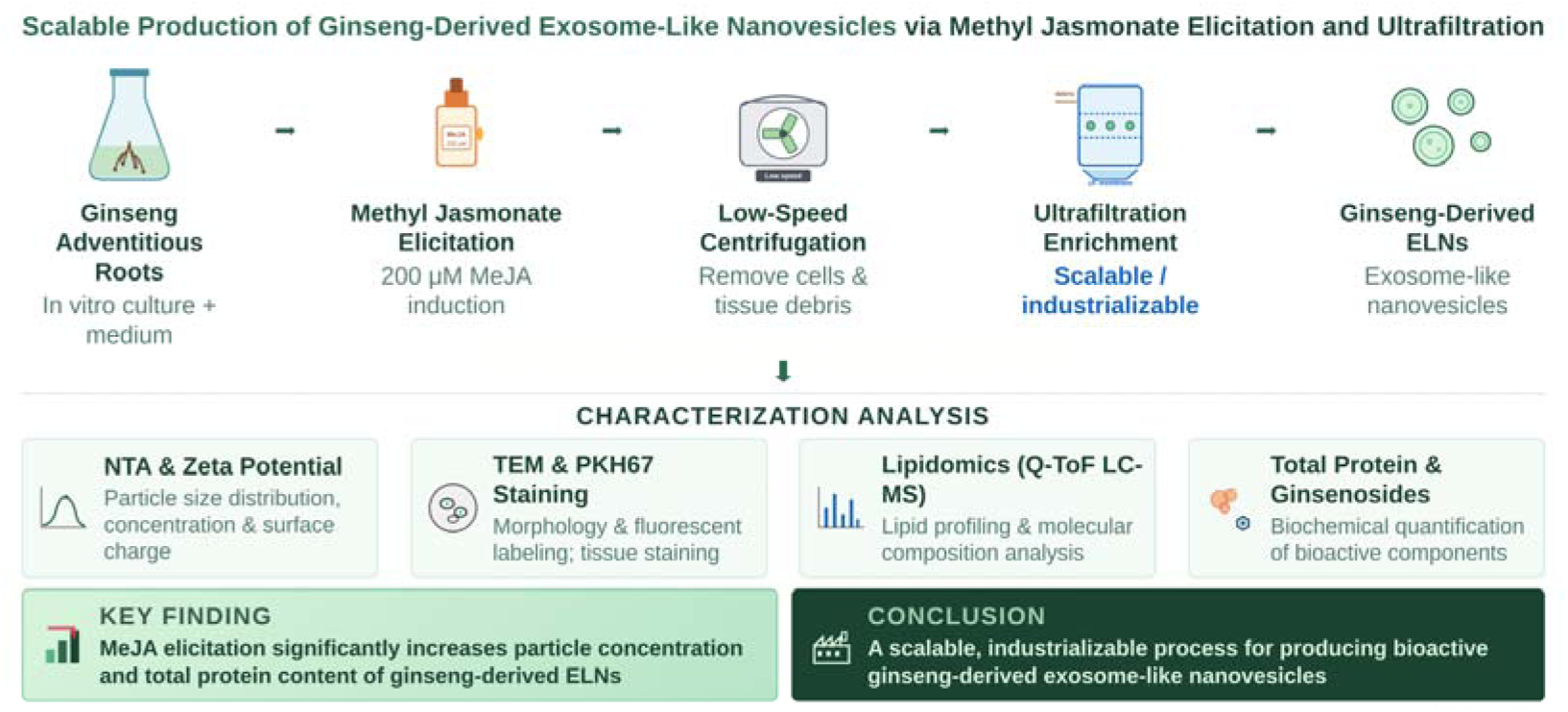

## Introduction

Ginseng (*Panax ginseng C. A. Meyer*) is a perennial herb of the Araliaceae family and the genus Panax. It has been known as the “king of herbs” and has a medicinal history of more than two thousand years in China. Modern pharmacological studies have shown that the active components of ginseng possess multiple pharmacological activities, including immunomodulation, antioxidant, antitumor, and neuroprotective effects ^[1[3]^. However, traditional research has mostly focused on ginseng extracts and small-molecule monomer components, while studies on nanoscale vesicles secreted by ginseng cells, namely exosome-like nanovesicles, remain relatively limited.

Plant-derived extracellular vesicles (pEVs) are nanoscale membrane vesicles secreted by plant cells into the extracellular space, with particle sizes ranging from 30 to 300 nm. They encapsulate bioactive molecules such as proteins, lipids, nucleic acids, and secondary metabolites ^[4[5]^. Recent studies have shown that pEVs can be taken up by mammalian cells and affect recipient cell functions through their carried miRNAs, proteins, and other substances, demonstrating a unique capacity for cross-kingdom communication^[6-[9]^. Compared with animal-derived exosomes, plant-derived exosome-like nanovesicles have broad sources, lower immunogenicity, and relatively low production costs, showing potential as natural bioactive delivery carriers ^[10[11]^. To date, pEVs have been successfully isolated from edible plants such as ginger, grape, lemon, and broccoli, and have been confirmed to possess biological functions including anti-inflammatory, antitumor, and intestinal homeostasis regulation effects ^[12[14]^. However, research on the isolation, extraction, and identification of exosome-like nanovesicles from medicinal plants such as ginseng is extremely limited.

Exosome-like nanovesicles actively secreted by ginseng adventitious roots into the culture medium during liquid culture are an ideal raw material for obtaining high-quality ginseng exosomes, because this source is mild, requires no tissue disruption and avoids intracellular impurity contamination. Currently, the core bottleneck restricting the industrialization of ginseng-derived exosome-like vesicles is the lack of separation technology suitable for low-concentration, large-volume culture supernatant systems that is simultaneously efficient, gentle, and amenable to scale-up. The commonly used isolation methods all have obvious shortcomings. Ultracentrifugation (dUC) is widely applied but suffers from low throughput, long processing time, and high equipment requirements, and high centrifugal force may cause structural damage to vesicles^[15]^. Polyethylene glycol (PEG) precipitation is simple to operate but yields products with low purity, and polymer residues interfere with subsequent functional evaluation^[16]^. Size exclusion chromatography (SEC) can better preserve vesicle integrity but has limited sample processing capacity^[17]^. None of these methods can readily achieve scale-up from laboratory to industrial production. To address these challenges, the present study constructed a novel isolation method for ginseng-derived exosome-like vesicles based on a magnetic stirred ultrafiltration cell. The morphology, physicochemical properties, and molecular composition of the isolated products were systematically identified and analyzed using transmission electron microscopy (TEM), nanoparticle tracking analysis (NTA), Zeta potential analysis, and Q-ToF LC-MS-based lipidomic analysis. The aim was to establish a standardized sample preparation system suitable for exosome-like vesicles derived from ginseng adventitious roots and their culture supernatant, and to develop a low-cost, membrane-fouling-resistant, and easily scalable pEV isolation process, providing technical support for the industrial production and application of ginseng-derived exosome-like nanovesicles.

## 1 Materials and Methods

### 1.1 Experimental Materials

#### 1.1.1 Ginseng Adventitious Roots

Ginseng (Korean ginseng and wild ginseng) adventitious roots in tissue culture and their culture supernatant (1/4 SH liquid medium), as well as corresponding samples induced with 200 μM methyl jasmonate (MeJA) for 10 days, were provided by Shandong Yanding Holding Group.

#### 1.1.2 Main Reagents

Phosphate-buffered saline (PBS, 0.01 M, pH 7.2) was prepared in our laboratory. SH basal salt medium with vitamins was purchased from Beijing Coolaber Technology Co., Ltd. Methyl jasmonate (MeJA, purity ≥ 95%) was purchased from Shanghai Macklin Biochemical Technology Co., Ltd. BeyoExo™ exosome labeling and tracing kit (PKH67) was purchased from Shanghai Beyotime Biotechnology Co., Ltd. BCA protein concentration assay kit (enhanced) was purchased from Shanghai Beyotime Biotechnology Co., Ltd. Nucleic acid extraction or purification reagent kit was purchased from Jiangsu Cowin Biotech Co., Ltd. SuperStar miRNA First-Strand cDNA Synthesis Kit (by tailing A) was purchased from Jiangsu Cowin Biotech Co., Ltd. SuperStar Universal SYBR Master Mix was purchased from Jiangsu Cowin Biotech Co., Ltd. Chromatographic grade dichloromethane and methanol were purchased from Merck (Germany). n-Butanol was purchased from Yantai Yuandong Fine Chemical Co., Ltd. Water-saturated n-butanol was prepared by mixing n-butanol and ultrapure water at a ratio of 1:1 (v/v), shaking thoroughly, allowing phase separation, and collecting the upper n-butanol phase. All water used in experiments was ultrapure water.

#### 1.1.3 Main Instruments

Multi-function blender (FB-110X15, Shanghai Litu Ultrahigh Pressure Equipment Co., Ltd.); low-speed centrifuge (GENIUS 6K-C, Changsha Xin’ao Instrument Co., Ltd.); filter (TC-20, Hangzhou Fuyang Huayi Instrument Co., Ltd.); 0.22 μm cellulose acetate membrane (Shanghai Xingya Purification Material Factory); magnetic stirrer (MYP19-2A, Shanghai Meiyinpu Instrument Manufacturing Co., Ltd.); stirred ultrafiltration cell (UFSC40001, Merck Life Science, Germany); 100 kDa ultrafiltration membrane (PBHK07610, Merck Life Science, Germany); UV-visible spectrophotometer (UV-6100, Shanghai Yuanxi Instrument Co., Ltd.); fluorescence microscope (BX53, Olympus, Japan); nanoparticle tracking analyzer (ZetaView, ParticleMetrix, Germany); nanoparticle size and zeta potential analyzer (Zetasizer Nano ZS90, Malvern, UK); transmission electron microscope (HT7700, Hitachi High-Tech, Japan); high-speed refrigerated centrifuge (FRESCO 21, Thermo Fisher Scientific, China); micro-volume spectrophotometer (SMA4000, Merinton Instrument, Beijing); gene amplification instrument (Thermo Fisher Scientific, China); high-resolution Q-ToF liquid chromatography-mass spectrometer (H-Class/Xevo G2-XS QTOF, Waters, USA).

### 1.2 Extraction of Ginseng Exosome-Like Nanovesicles

Exosome-like nanovesicles were isolated from ginseng adventitious root tissue and culture supernatant, respectively.

(1) Pretreatment of adventitious root samples Fresh sterile adventitious roots (150 g) were rinsed with sterile ultrapure water to remove residual medium, cut into pieces, and mixed with pre-cooled sterile PBS (4°C, pH 7.2-7.4) at a material-to-liquid ratio of 1:5. The tissue was homogenized using a high-speed blender to prepare the crude tissue extract.
(2) Pretreatment of culture supernatant Culture supernatant (2.5 L) from adventitious roots was filtered through a 200-mesh sieve to remove large particulate debris.
(3) Low-speed centrifugation and filtration The crude extract and filtrate were centrifuged at 3500 r/min for 30 min. The supernatant was collected and filtered through a 0.22 μm cellulose acetate membrane to remove particulate impurities and microorganisms.
(4) Concentration and diafiltration using a magnetic stirred ultrafiltration cell The filtrate was transferred to a stirred ultrafiltration cell equipped with a 100 kDa PES ultrafiltration membrane. The working pressure was 0.15-0.25 MPa, and the stirring speed was 80-120 r/min. The feed was concentrated to 1/8-1/10 of the original volume, followed by two rounds of diafiltration with an equal volume of sterile PBS to remove free small molecules. The membrane-retained fraction was collected as the vesicle extract.
(5) Sample processing and storage The vesicle extract was filtered through a 0.22 μm PES sterile membrane. Samples were stored at 4°C for short-term use and at −80°C for long-term storage in aliquots, avoiding repeated freeze-thaw cycles.

### 1.3 Characterization and Physicochemical Analysis of Exosome-Like Nanovesicles

Transmission electron microscopy (TEM), nanoparticle tracking analysis (NTA), Zeta potential analysis, and lipid composition analysis were used to comprehensively characterize the morphology, particle size distribution, particle concentration, protein concentration, and lipid composition of the extracted ginseng-derived exosome-like samples.

#### 1.3.1 TEM Morphology Observation

A 20 μL aliquot of vesicle sample was dropped onto a copper grid and allowed to adsorb for 10 min. Excess liquid was removed with filter paper. The sample was negatively stained with phosphotungstic acid for 30 s, dried under an infrared lamp, and observed by transmission electron microscopy to record the bilayer membrane morphology.

#### 1.3.2 NTA and Zeta Potential Analysis

Samples were appropriately diluted with sterile water. NTA was used to detect particle size distribution and particle concentration, with three biological replicates per group and a single acquisition time of 60 s. Diluted samples were used for Zeta potential measurement at 25°C with a 120 s equilibration time, and three parallel measurements were performed per group. Results are expressed as mean ± standard deviation (mean ± SD).

#### 1.3.3 PKH67 Fluorescence Labeling

A 50 μL aliquot of sample was labeled with the BeyoExo™ PKH67 exosome labeling and tracing kit according to the manufacturer’s instructions. After labeling, samples were mounted and observed under a fluorescence microscope for green fluorescence signal distribution.

#### 1.3.4 Paraffin Section Staining of Ginseng Adventitious Root Tissue

Ginseng adventitious root tissue was fixed in FAA fixative, dehydrated through a graded ethanol series, cleared with xylene, embedded in paraffin, and sectioned at 4-6 μm thickness. After deparaffinization and rehydration, sections were stained with DAPI, neutral red, and PKH67, respectively. After dehydration and clearing, sections were mounted with neutral balsam and observed under a fluorescence microscope for fluorescence signal distribution within the tissue.

#### 1.3.5 BCA Total Protein Content Determination

The enhanced BCA protein assay kit was used. Absorbance was measured at 562 nm, and total protein concentration of vesicle samples was calculated from the standard curve, with three replicates per group.

#### 1.3.6 Lipid Extraction and Q-ToF LC-MS Lipidomic Analysis

Referring to the method reported by Folch et al.^[18]^with minor modifications, samples were extracted with a methanol-water-dichloromethane system. The lower organic phase was collected by centrifugation, washed with pure water, and dried with anhydrous sodium sulfate. After solvent evaporation, samples were reconstituted in methanol-isopropanol (1:1, v/v) for LC-MS analysis.

Liquid chromatography was performed on an ACQUITY UPLC® CSH C18 column with gradient elution. Mass spectrometric data were acquired in both positive and negative ion modes with an ESI source, and the mass scan range was m/z 50-1200 Da. Progenesis QI software was used to process mass spectrometric data, and lipid molecules were qualitatively identified by comparison with LipidBlast, LMSD, and HMDB databases.

#### 1.3.7 Total Saponin Content Determination

Referring to the national standard method GB/T 22248-2008, a modified vanillin-sulfuric acid colorimetric method was used. Vesicle samples were extracted with water-saturated n-butanol, evaporated to dryness, and reconstituted with methanol. Then 8% vanillin-ethanol and 72% sulfuric acid were added, and the mixture was reacted in a 60°C water bath for 10 min, cooled in an ice bath, and absorbance was measured at 560 nm. Total saponin content was calculated using a standard curve established with ginsenoside Re.

### 1.4 miRNA Extraction, Reverse Transcription, and RT-qPCR Analysis

Eight experimental groups were established: exosome-like vesicles derived from culture supernatant and adventitious roots of Korean ginseng and wild ginseng, divided into non-induced control (C) and 200 μM MeJA-induced (I) groups, with three biological replicates per group.

Vesicular miRNA was extracted, and RNA purity and concentration were determined. miRNA reverse transcription was performed using a poly(A) tailing kit to obtain cDNA. The qPCR reaction system was 20 μL, with U6 snRNA as the internal reference gene, and three technical replicate wells were set for each sample. After the reaction, melting curve analysis was performed to verify amplification specificity. The 2^-ΔΔCt method was used to calculate relative miRNA expression. Differentially expressed miRNAs were screened using |log FC| ≥ 1 and *p* < 0.05 as criteria. Principal component analysis (PCA), Z-score standardization, and clustering heatmap visualization were performed. All experimental results are expressed as mean ± SD, and *p* < 0.05 was considered statistically significant.

## 2 Results

### 2.1 Extraction of Ginseng-Derived Exosome-Like Nanovesicles

Based on the extraction procedure described in Materials and Methods, exosome-like nanovesicles were obtained from ginseng adventitious roots and their culture supernatant, respectively. Samples from different sources were processed using the same unified procedure, and the obtained samples were used for subsequent analysis of morphology, particle size, particle concentration, Zeta potential, and biochemical composition.

### 2.2 Phenotypic Analysis of Ginseng Exosome-Like Nanovesicles

TEM analysis showed that at 50k magnification, cup-shaped, saucer-shaped, or circular bilayer vesicle structures were observed in both Korean ginseng and wild ginseng adventitious roots and their culture supernatant before and after methyl jasmonate induction (Figure 1), consistent with typical plant-derived exosome-like nanovesicle structures.

**Fig. 1.**
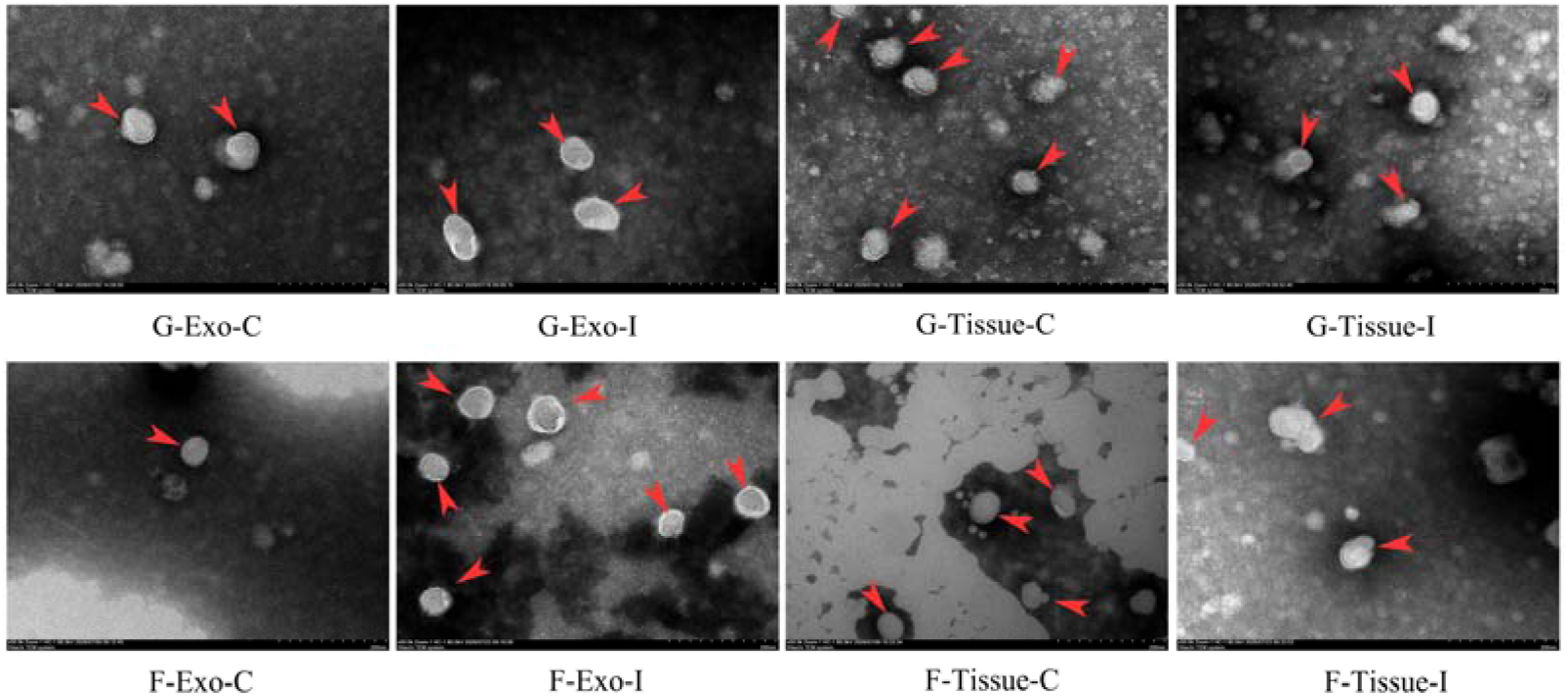
TEM image of ginseng exosome like nanovesicles. Note: G, Korean ginseng; F, wild ginseng; Exo, extracellular vesicle/exosome; Tissue, tissue-derived EV; C, control (before induction/untreated); I, induce. Scale bar: 200 nm. Red arrows indicate typical cup-shaped, saucer-shaped, or circular bilayer vesicle structures.

PKH67 fluorescence staining results (Figure 2) showed that at 1000 magnification, all eight sample groups exhibited uniformly distributed and dense green fluorescence labeling. In contrast, the PBS buffer control group (CK) showed no uniformly distributed or dense green fluorescence signals, indicating the presence of lipid membrane vesicles that can be labeled by PKH67 in all samples.

**Fig. 2.**
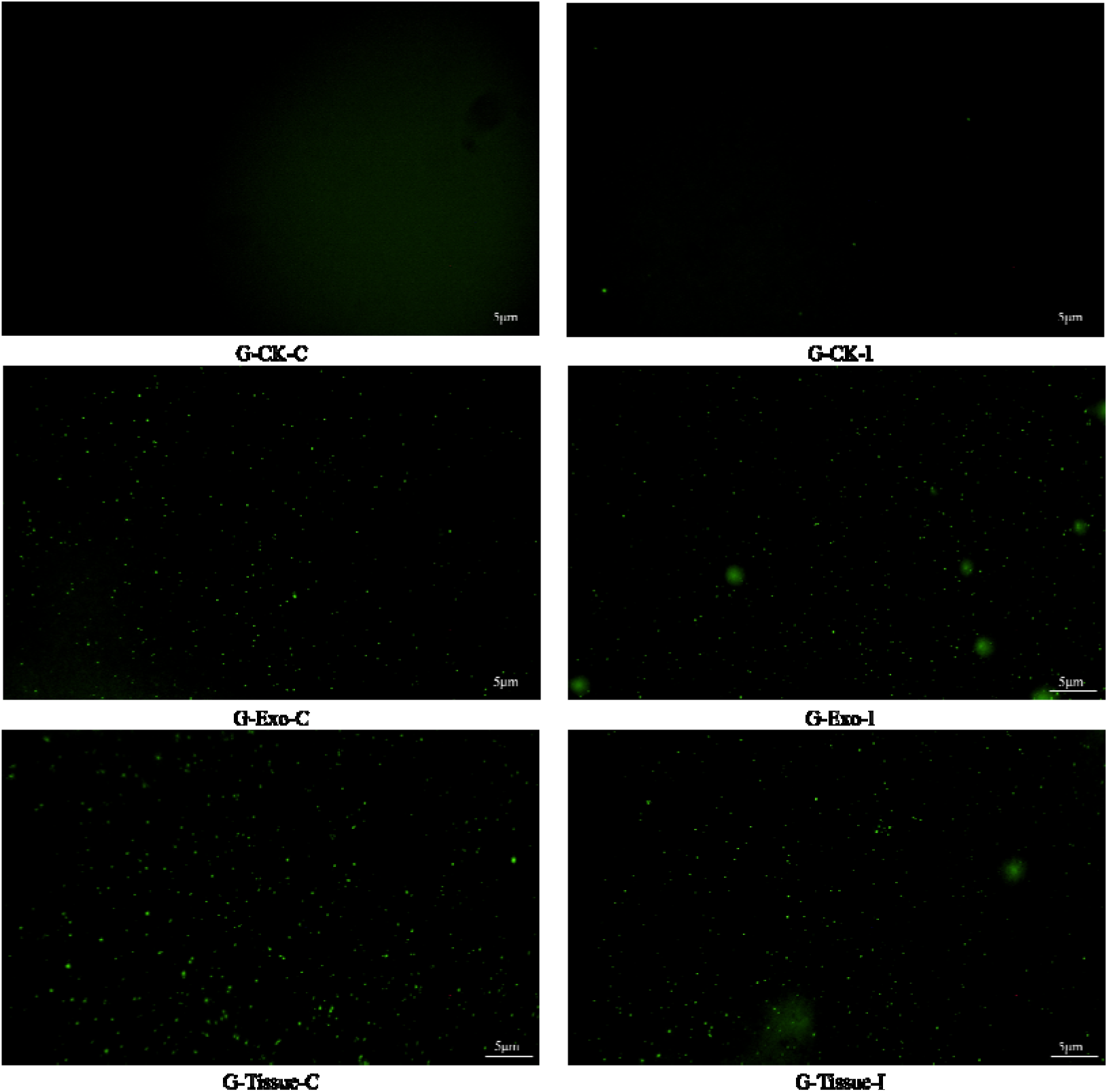
Fluorescence staining of PKH67 in ginseng exosome like nanovesicles (Partial). Note: G, Korean ginseng; Exo, extracellular vesicle/exosome; Tissue, tissue-derived EV; C, control (before induction/untreated); I, induce; CK, PBS buffer control group.

NTA analysis results (Table 1) showed that the average particle sizes of the eight ginseng-derived exosome-like nanovesicle samples ranged from 77.333 nm to 171.733 nm, and the particle concentration ranged from 2.1E+8 /mL to 7.6E+10 /mL. Compared with non-induced samples, the particle concentration of exosome-like nanovesicles from samples induced with methyl jasmonate was increased. Combined with the particle size distribution plots (Figure 3), all samples showed particle sizes mainly distributed in the range of 70 nm to 300 nm, consistent with the typical particle size characteristics of plant-derived exosome-like nanovesicles. Meanwhile, the dispersity of most samples was below 0.410, with only a few samples showing a dispersity of 0.505, indicating that the obtained nanovesicles had good dispersity and uniform distribution.

**Fig. 3.**
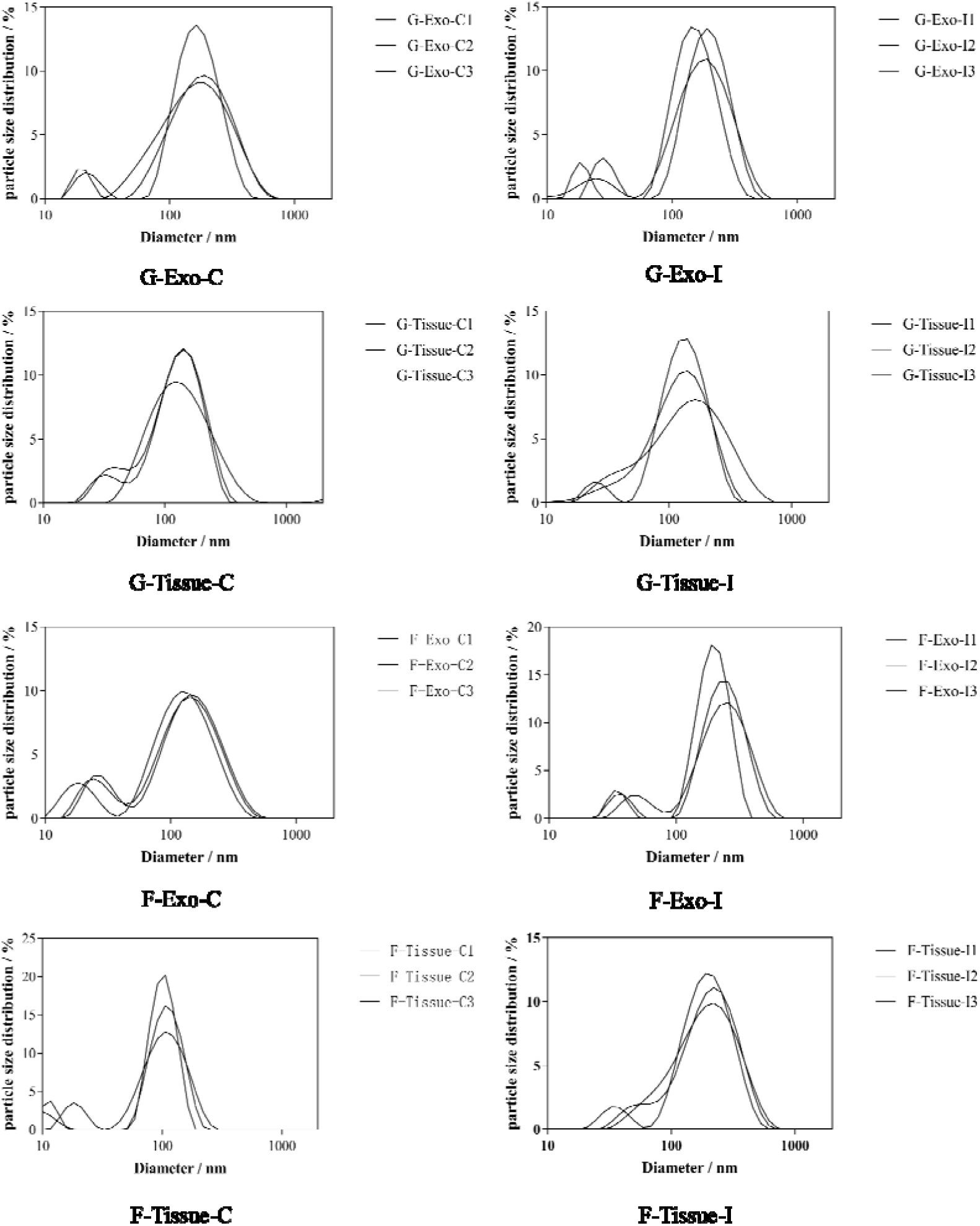
Particle size distribution map of ginseng derived exosome like nanovesicles. Note: G, Korean ginseng; F, wild ginseng ; Exo, extracellular vesicles/exosomes; Tissue, Tissue derived EV; C, Control (pre induction/untreated); I, Induce.

**Table 1.** Analysis of NTA and Zeta Potential of Extracellular vesicle like Nanovesicles.

| Sample | average particle size(nm) | dispersity ( PdI ) | Particle concentration/mL | Zeta potential (mV) |
| --- | --- | --- | --- | --- |
| G-Exo-C | 121.633±4.751 | 0.375±0.080 | 4.7E+8±8.2E+7 | -16.567±1.167 |
| G-Tissue-C | 111.767±2.786 | 0.382±0.003 | 2.3E+10±3.7E+9 | -12.767±0.403 |
| G-Exo-I | 120.500±7.201 | 0.380±0.050 | 4.9E+8±5.5E+7 | -10.880±1.523 |
| G-Tissue-I | 111.600±9.706 | 0.366±0.029 | 7.6E+10±1.6E+10 | -14.233±0.170 |
| F-Exo-C | 77.333±2.855 | 0.505±0.017 | 4.2E+8±7.1E+7 | -1.830±0.467 |
| F-Tissue-C | 91.510±10.847 | 0.410±0.043 | 2.1E+8±3.6E+7 | -10.017±0.997 |
| F-Exo-I | 171.733±8.599 | 0.362±0.062 | 6.9E+8±2.4E+7 | -4.530±0.738 |
| F-Tissue-I | 151.967±8.599 | 0.272±0.002 | 6.8E+9±2.6E+9 | -11.800±0.283 |

Zeta potential analysis results (Table 1) showed that the Zeta potential of the eight ginseng-derived exosome-like nanovesicle samples ranged from −1.830 to - 16.567 mV. All samples exhibited negative charge, indicating that ginseng-derived exosome-like nanovesicles have negatively charged surface characteristics in PBS buffer system. Differences in Zeta potential among different samples suggest that surface charge properties may be influenced by sample source and processing conditions.

To investigate the origin of ginseng-derived exosome-like nanovesicles, paraffin sections of ginseng adventitious root tissue were stained with PKH67, neutral red, and DAPI. The results (Figure 4) showed that at 200 magnification, cell nuclei were clearly observed, and neutral red and PKH67 staining results largely overlapped, preliminarily suggesting that some ginseng-derived exosome-like nanovesicles may be associated with vacuoles.

**Fig. 4.**
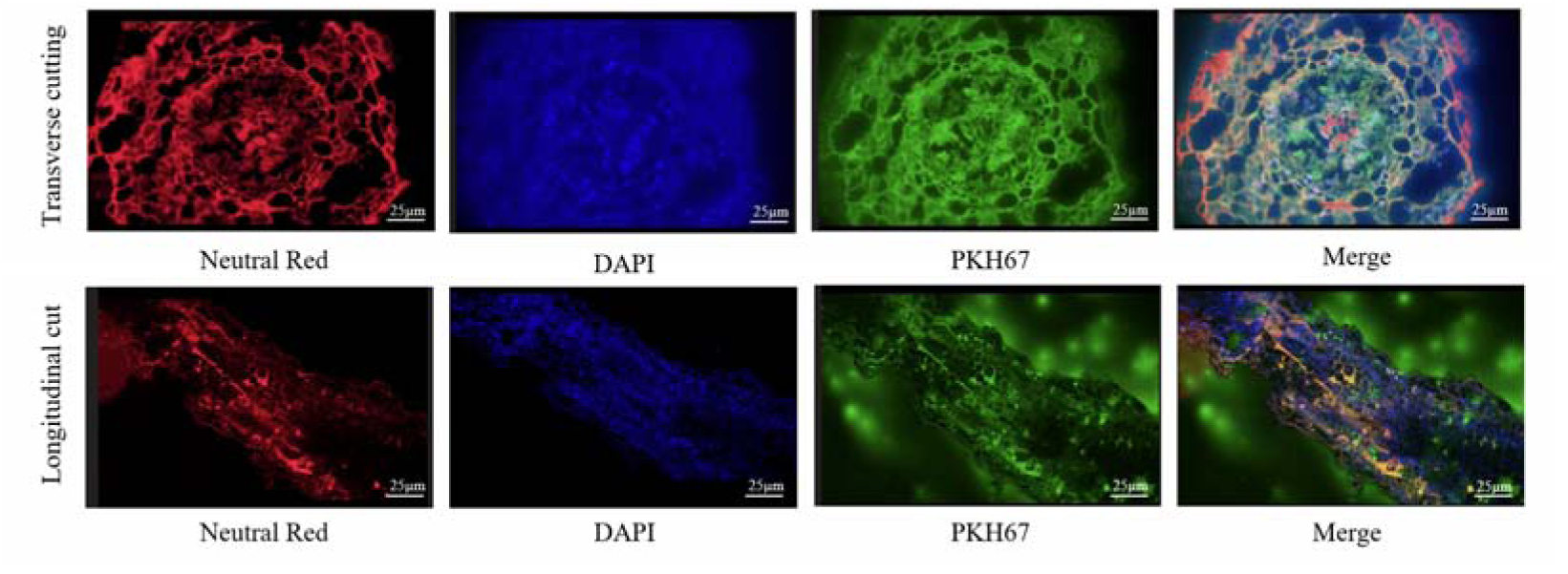
Staining of paraffin sections of ginseng adventitious root tissue.

### 2.3 Physicochemical Properties of Ginseng-Derived Exosome-Like Nanovesicles

#### 2.3.1 Protein Content Determination

BCA protein assay results (Figure 5) showed that the total protein concentration of the eight sample groups ranged from 0.685 to 8.185 mg/mL. The protein concentration standard curve is shown in Figure 5A, with a regression equation of Y = 2.017X -0.002783 and a coefficient of determination R² = 0.9978, indicating good linearity and suitability for determining total protein concentration in samples. Comparison among samples from different sources revealed that the total protein concentration of exosome-like nanovesicles from culture supernatant was generally higher than that from ginseng adventitious roots. Compared with the non-induced group, the total protein concentration of exosome-like nanovesicles from both ginseng adventitious roots and culture supernatant was increased after methyl jasmonate (MeJA) induction (Figure 5B).

**Fig. 5.**
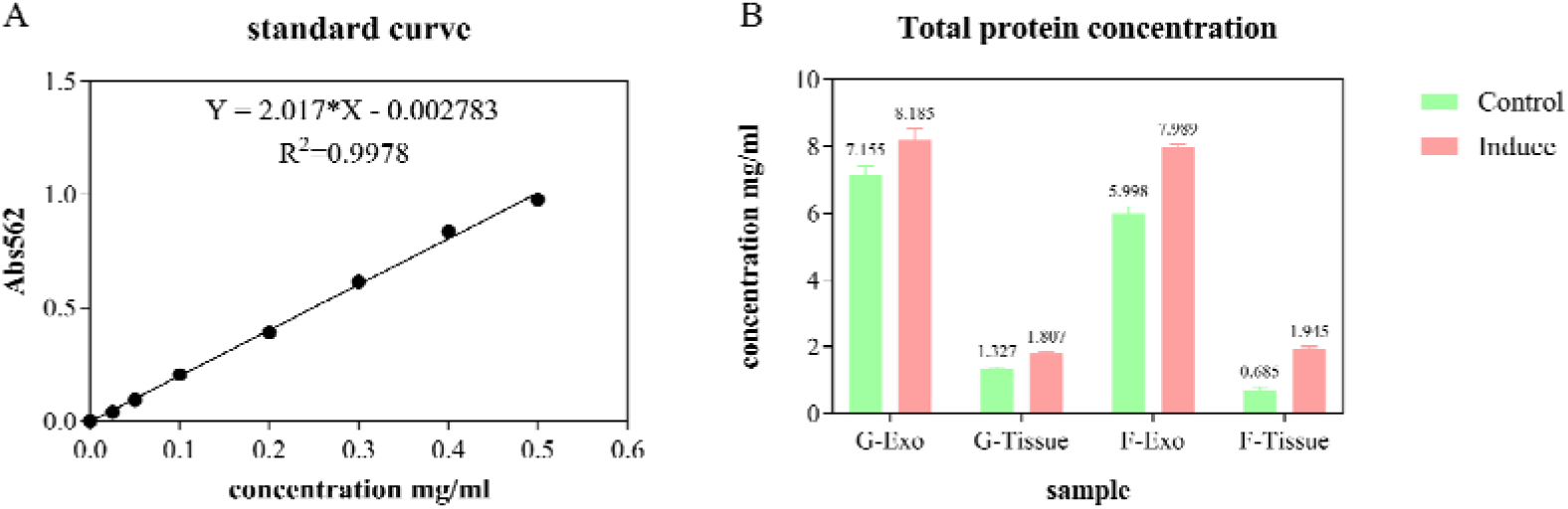
Total protein concentration using BCA method. Note: G, Korean ginseng; F, wild ginseng; Exo, Extracellular vesicles/exosomes; Tissue, Tissue derived EV; C, Control (pre induction/untreated); I, Induce.

#### 2.3.2 Lipidomic Analysis of Ginseng-Derived Exosomes

Lipid composition analysis of exosome-like nanovesicles from wild ginseng adventitious roots and their culture supernatant before induction showed (partial results in Tables 2 and 3) that multiple lipid components were detected in ginseng-derived exosome-like nanovesicles, mainly including triglycerides (TG), diglycerides (DG), phosphatidic acid (PA), phosphatidylcholine (PC), Cer/Sphingoid, and phosphatidylethanolamine (PE). A total of 165 peaks and 6,214 lipid entries were detected in exosome-like nanovesicles from culture supernatant, while 65 peaks and 552 lipid entries were detected in those from adventitious roots, lower than those from culture supernatant (Supplementary Tables S2 and S3).

**Table 2.**
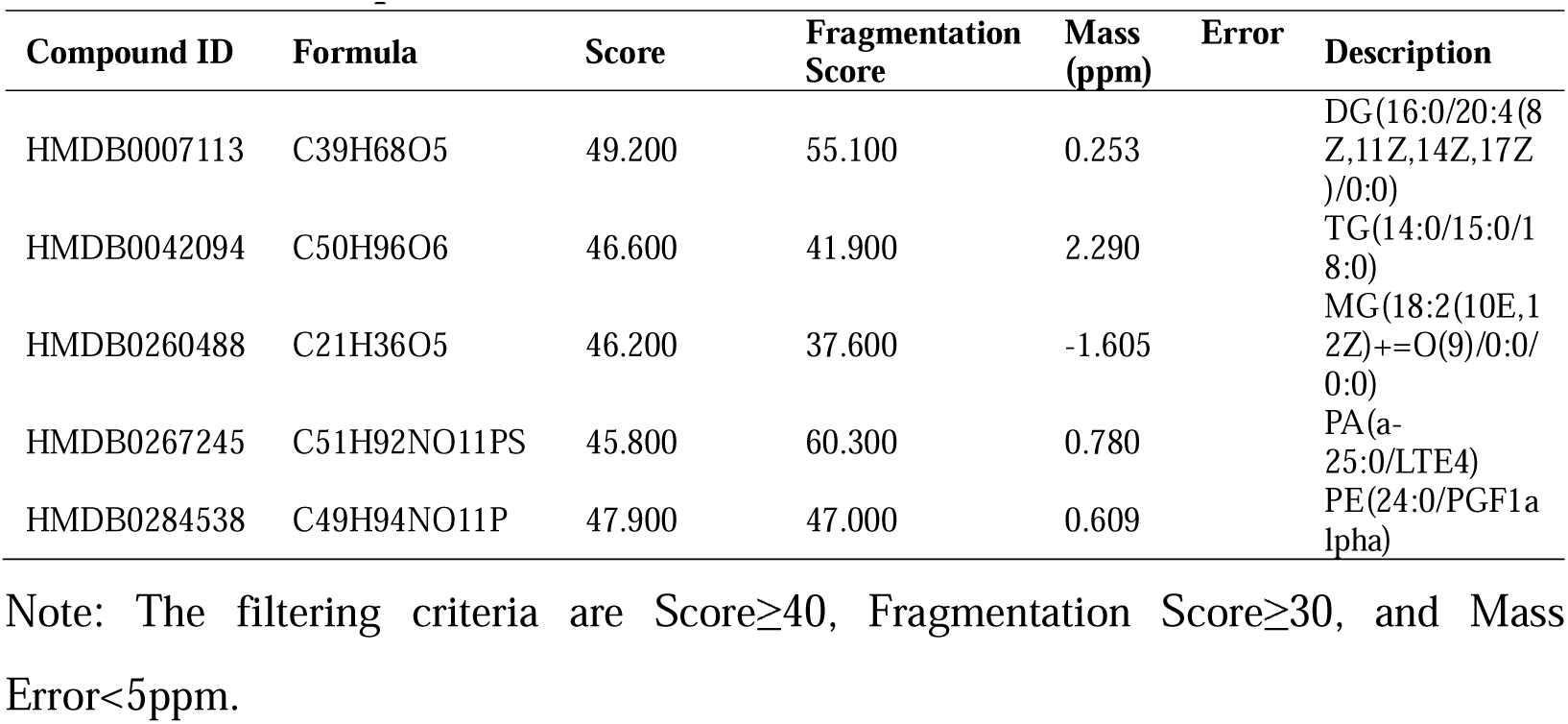
Analysis of lipid content in ginseng exosome like nanovesicles derived from the culture supernatant (Partial)

**Table 3.**
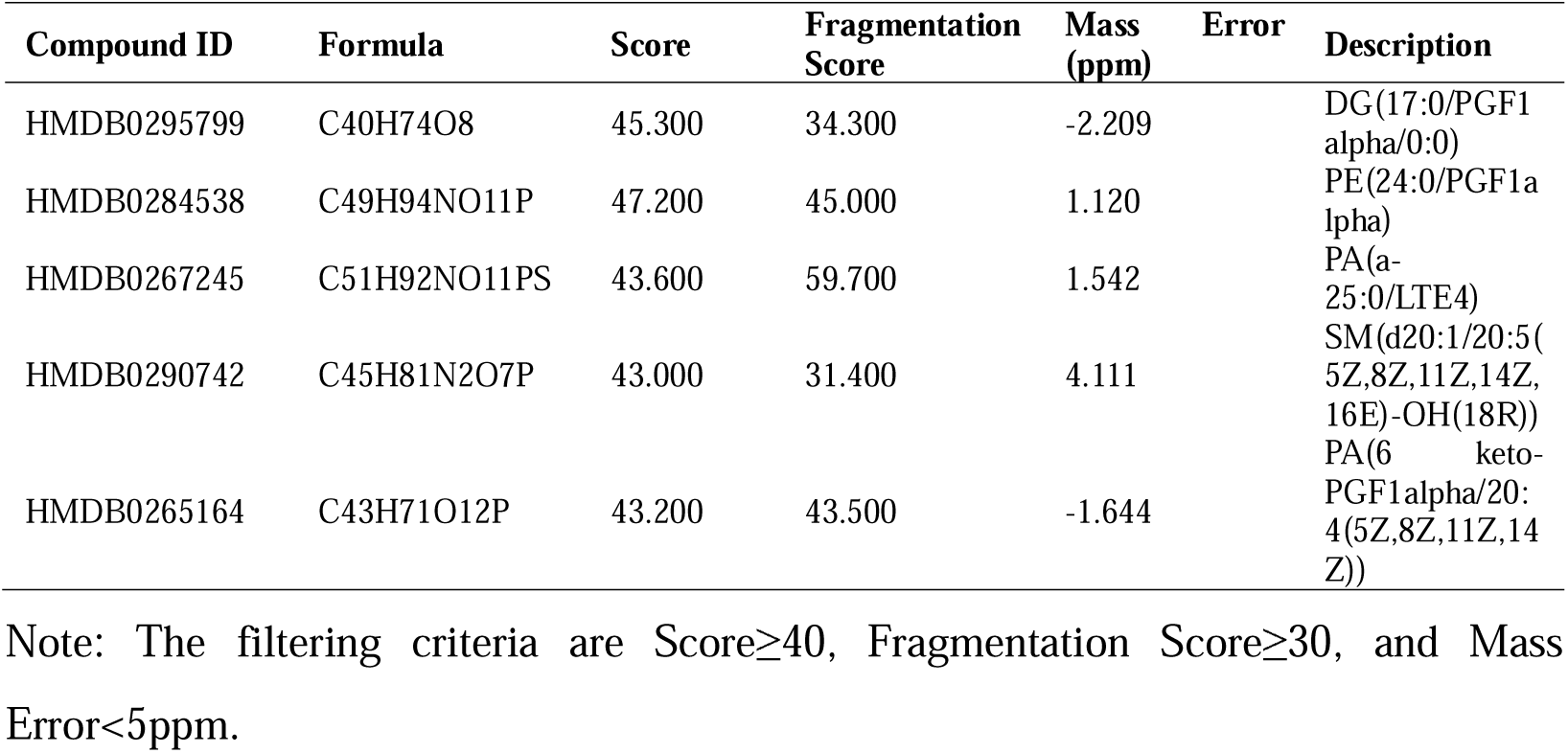
Lipid analysis of extracellular vesicle like nano vesicles derived from tissue cultured adventitious roots of ginseng (Partial)

#### 2.3.3 Total Saponin Content Determination in Ginseng Exosomes

Referring to the national standard method GB/T 22248-2008, the total saponin concentration of eight ginseng-derived exosome-like nanovesicle samples and four ginseng adventitious root samples was determined. The results are shown in Figure 7. The total saponin concentration of the eight exosome-like nanovesicle samples ranged from 0.010% to 0.079%, while that of the four ginseng adventitious root samples ranged from 1.764% to 7.622%. The saponin concentration standard curve is shown in Figure 6A, with a regression equation of Y = 0.02037X - 0.01884 and a coefficient of determination R² = 0.9903, indicating good linearity and suitability for determining total saponin concentration in samples.

**Fig. 6.**
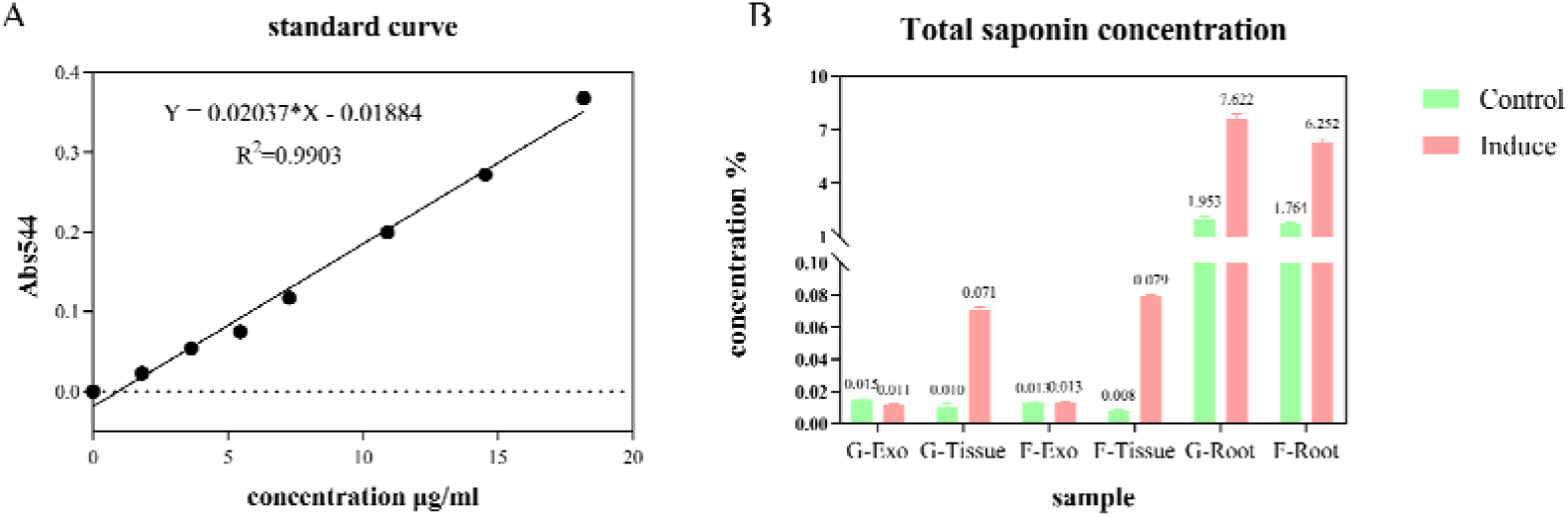
Total saponin concentration. Note: G, Korean ginseng; F, wild ginseng; Exo, Extracellular vesicles/exosomes; Tissue, Tissue derived EV; Root, Ginseng fibrous root; C, Control (pre induction/untreated); I, Induce.

**Fig. 7.**
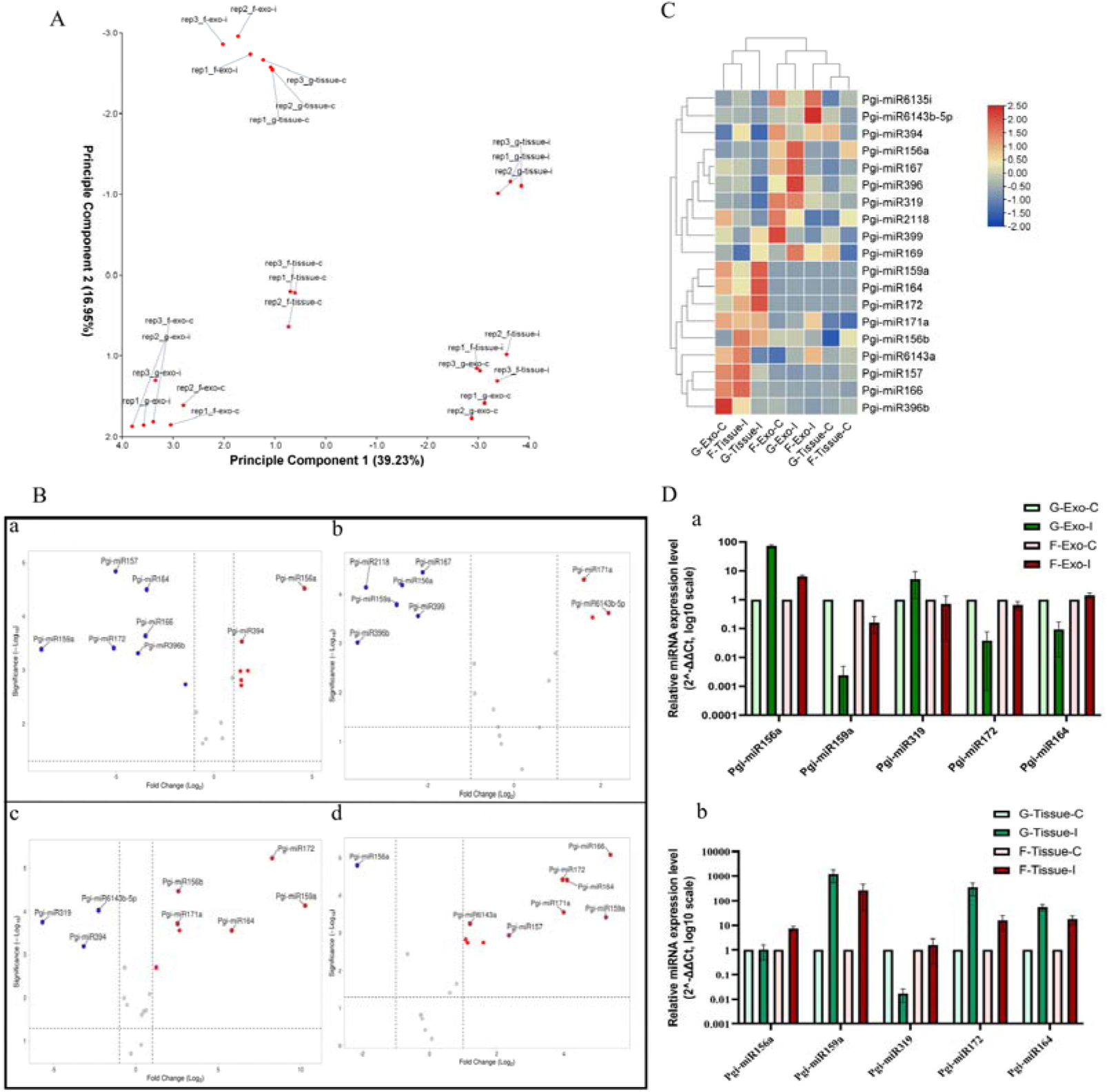
miRNA fluorescence quantitative PCR. Note: **A**, Principal Component Analysis (PCA) of candidate miRNAs in different experimental groups; **B**, Volcanic diagram of differential expression of candidate miRNAs before and after induction of samples from different sources, a: G-Exo-I vs G-Exo-C (induced by exosomes derived from ginseng culture medium vs before induction), b: F-Exo-I vs F-Exo-C (induced by exosomes derived from wild ginseng culture medium vs before induction), c: G-Tissue-I vs G-Tissue-C (induced by adventitious root tissue cultured samples of ginseng vs before induction), d: F-Tissue-I vs F-Tissue-C (induced by adventitious root tissue cultured samples of wild ginseng vs before induction). The screening criteria are | log FC | ≥ 1, *p*<0.05; **C**, Z-score clustering heatmap of candidate miRNA expression patterns in different experimental groups; **D**, Relative expression levels of differentially expressed miRNAs before and after induction in samples from different sources, a: samples from culture medium sources; b: Tissue culture samples from adventitious root sources.

Comparison among samples from different sources revealed that the total saponin content of ginseng-derived exosome-like nanovesicles was much lower than that of ginseng adventitious roots. Before induction, the total saponin concentration of exosome-like nanovesicles from culture supernatant was generally higher than that from ginseng adventitious roots, but after induction, the total saponin concentration of exosome-like nanovesicles from culture supernatant was lower than that from ginseng adventitious roots. Compared with the non-induced group, after methyl jasmonate (MeJA) induction, the total saponin concentration of both ginseng adventitious roots and ginseng adventitious root-derived exosome-like nanovesicles was significantly increased (Figure 6B), while the total saponin concentration of culture supernatant-derived exosome-like nanovesicles showed no differential change.

### 2.4 Ginseng miRNA Fluorescence Quantitative PCR

Based on candidate miRNA expression data, PCA analysis (Figure 7A) showed that the first two principal components cumulatively explained 56.18% of sample variation. Biological replicates in each group clustered together in PCA space, and samples from different sources and treatment conditions showed some degree of spatial separation. Among them, exosomes derived from Korean ginseng and wild ginseng adventitious roots and their culture supernatant showed different distribution characteristics, indicating that sample source and MeJA induction treatment were associated with changes in the overall expression pattern of candidate miRNAs.

Four volcano plots (Figure 7B) compared miRNA expression differences before and after induction in various samples. Induction treatment significantly altered the miRNA expression pattern of exosomes from Korean ginseng culture supernatant. Exosomes from wild ginseng culture supernatant exhibited a miRNA response pattern distinct from that of Korean ginseng exosomes. Both Korean ginseng and wild ginseng adventitious root samples showed significant miRNA upregulation and downregulation after induction, suggesting that induction treatment can cause changes in the candidate miRNA expression profile.

Z-score standardization and hierarchical clustering analysis (Figure 7C) revealed that candidate miRNAs showed significant differentiated expression patterns across the eight experimental groups, with some miRNAs enriched in culture supernatant exosomes and others preferring adventitious root samples. Multiple miRNAs formed clustering modules with similar expression trends, presumably influenced by common regulatory mechanisms. Overall, sample source and MeJA induction treatment were both associated with changes in candidate miRNA expression patterns.

As shown in Figure 7D, each miRNA exhibited a distinct response pattern to induction treatment. The expression changes of Pgi-miR156a (plant development-related), Pgi-miR159a (hormone response-related), Pgi-miR319 (secondary metabolism-related), Pgi-miR172 (plant development-related), and Pgi-miR164 (plant development-related) combined induction effects with sample source specificity. These quantitative results were consistent with the conclusions from PCA, volcano plots, and clustering heatmaps, indicating that candidate miRNAs exhibit differentiated expression patterns under different sources and MeJA induction conditions.

## 3 Discussion

In recent years, plant-derived exosome-like nanovesicles have gradually become a research hotspot as natural active substance delivery carriers due to their broad material sources, good biocompatibility, and potential low immunogenicity. Effective extraction and characterization of plant-derived exosome-like nanovesicles is an important basis for conducting research on their biological activities and applications. Jae et al.^[19]^ extracted and identified exosome-like nanovesicles from cabbage and red cabbage, and the results showed that the products were spherical with an average particle size of approximately 100 nm and Zeta potentials of −14.8 mV and −15.2 mV. Sriwastva et al.^[20]^ isolated mulberry bark-derived exosome-like nanovesicles, and TEM observation showed that they had a bilayer membrane structure, with NTA-determined average particle diameter of 151.3 ± 45.4 nm. Zhao et al.^[21]^ extracted plant-derived exosomes from 14 plants including blueberry, yam, bitter melon, and peach, and performed particle size distribution and potential analysis on blueberry exosome-like nanovesicles, which showed particle sizes ranging from 150 to 250 nm and a Zeta potential of −2.52 mV. The ginseng-derived exosome-like nanovesicles obtained in this study were basically consistent with the above research results in terms of morphology and particle size, further indicating that the established extraction method can obtain ginseng-derived exosome-like nanovesicles with typical exosome characteristics.

Plant-derived exosome-like nanovesicles have a lipid bilayer membrane structure, and lipophilic fluorescent dyes such as PKH67 can fluorescently label lipid vesicles through interaction with membrane lipids^[22]^. In this study, PKH67 green fluorescent dye was used to label ginseng-derived exosome-like nanovesicles, and relatively uniform green fluorescence signals were observed under a fluorescence microscope, while no obvious green fluorescence signals were observed in the PBS control group, indicating the presence of lipid vesicles that can be labeled by PKH67 in the obtained samples. Cho et al.^[23]^ analyzed the lipid composition of ginseng-derived extracellular vesicles and found that they contained lipid components such as phosphatidylcholine (PC), phosphatidylethanolamine (PE), diacylglycerol (DG), triacylglycerol (TG), monoacylglycerol (MG), and phosphatidic acid (PA). Liu^[24]^ found that Arabidopsis-derived extracellular vesicles also contain various lipid components including sphingolipids, phospholipids, and triglycerides. The lipidomic analysis results of this study are basically consistent with the above studies, further supporting the characteristics of the obtained ginseng-derived exosome-like nanovesicles from the perspective of lipid composition.

It is now generally believed that plant-derived exosome-like nanovesicles have three origin pathways: multivesicular body pathway, exocyst-positive organelle pathway, and vacuolar pathway^[25]^. Meanwhile, neutral red staining can cause vacuoles in ginseng cells to emit red fluorescence signals^[26]^. Combined with the tissue section staining results of ginseng adventitious roots in this study, neutral red fluorescence signals largely overlapped with PKH67 fluorescence signals, preliminarily suggesting that some ginseng-derived exosome-like nanovesicles may originate from the vacuolar pathway in ginseng cells.

Previous studies have shown that plant-derived exosome-like nanovesicles can carry bioactive molecules such as miRNAs and may participate in cell-to-cell information exchange and plant-animal interactions, with potential roles in regulating immune responses and metabolic homeostasis^[27]^. Methyl jasmonate (MeJA), as an important inducer of the plant jasmonic acid signaling pathway, can participate in plant growth and development, defense responses, and secondary metabolism regulation^[28]^. On the basis of establishing the extraction method for ginseng-derived exosome-like nanovesicles, this study used 200 μM MeJA to induce ginseng adventitious roots and compared changes in exosome-like nanovesicle concentration and miRNA expression levels before and after induction. NTA and total protein concentration determination results showed that after MeJA induction, the particle concentration and total protein concentration of ginseng-derived exosome-like nanovesicles were increased, indicating that MeJA treatment may promote the production or release of exosome-like nanovesicles from ginseng adventitious roots. Total saponin concentration determination results showed that after MeJA induction, the saponin content of ginseng adventitious root-derived exosome-like nanovesicles was greatly increased, while the saponin content of ginseng culture supernatant-derived exosome-like nanovesicles showed no specific change, indicating that MeJA treatment may promote the saponin content of ginseng adventitious root-derived exosome-like nanovesicles but has no obvious effect on culture supernatant-derived exosome-like nanovesicles. Meanwhile, real-time fluorescence quantitative PCR results showed that the expression levels of miRNAs such as Pgi-miR156a, Pgi-miR159a, and Pgi-miR319 changed before and after induction, which are respectively related to plant development, hormone response, and secondary metabolism. These results suggest that MeJA induction may affect the yield and miRNA molecular composition of ginseng adventitious root-derived exosome-like nanovesicles.

The potential mechanism by which MeJA induction increases exosome-like nanovesicle concentration may be related to enhanced plant defense and secondary metabolism responses. As an important plant signaling molecule, MeJA can activate multiple defense and secondary metabolism-related signaling pathways and may affect the cell membrane system and vesicle trafficking processes, thereby promoting the formation, accumulation, and release of extracellular vesicles. Therefore, the increased exosome-like nanovesicle concentration and total protein concentration observed in this study may be a comprehensive result of plant cell physiological responses induced by MeJA. MeJA can activate the ginsenoside biosynthesis pathway in ginseng adventitious root cells^[29]^, promoting the encapsulation and loading of saponin molecules by intracellular vesicles. Therefore, the saponin content of adventitious root-derived exosome-like nanovesicles was significantly increased. However, MeJA did not significantly promote the release of these high-saponin-loaded vesicles into the extracellular culture supernatant. Combined with the dilution of the culture supernatant system, easy degradation of vesicles, and complex vesicle population composition, the final result was no obvious specific change in saponin content of culture supernatant-derived vesicles. However, the present study still lacks direct evidence at the gene expression or protein level related to vesicle formation, exocytosis, and membrane vesicle trafficking, so the above mechanism still needs to be verified through further experiments.

This study still has certain limitations. The absolute Zeta potential values of some samples were relatively low, with the lowest value being −1.830 mV. Zeta potential is influenced by multiple factors such as dispersion medium ionic strength, pH, and vesicle surface composition. The relatively high ionic strength of PBS may compress the electric double layer on the vesicle surface, thereby affecting Zeta potential measurement results^[30]^. Therefore, subsequent studies can further optimize Zeta potential detection conditions, for example, by using a suitable low ionic strength buffer system and standardizing sample concentration and detection conditions to improve the accuracy of Zeta potential comparison among different samples. At the same time, gene expression analysis related to vesicle formation and extracellular release can be combined to further verify the potential mechanism by which MeJA promotes the production or release of ginseng-derived exosome-like nanovesicles at the molecular level.

## 4 Conclusion

This study established a method for extracting ginseng-derived exosome-like nanovesicles based on low-speed centrifugation, microfiltration, and 100 kDa ultrafiltration membrane combination, and successfully obtained exosome-like nanovesicles from ginseng adventitious roots and their culture supernatant. Through comprehensive characterization of the obtained samples by transmission electron microscopy (TEM), nanoparticle tracking analysis (NTA), Zeta potential analysis, and lipid composition analysis, the results showed that the obtained samples had typical bilayer membrane structures and corresponding nanoscale particle size distribution characteristics, and the lipid composition was also consistent with related studies on plant-derived extracellular vesicles, indicating that this method can be used for the preparation and subsequent research of ginseng-derived exosome-like nanovesicles.

The study further found that both ginseng adventitious roots and their culture supernatant can serve as sample sources for exosome-like nanovesicles, and samples from different sources showed certain differences in particle concentration, total protein, and total saponin levels. After induction with 200 μM methyl jasmonate for 10 days, the particle concentration and total protein levels of some source samples changed, and the total saponin concentration of adventitious root-derived exosome-like nanovesicles was significantly increased. Meanwhile, the expression patterns of candidate miRNAs such as Pgi-miR156a, Pgi-miR159a, Pgi-miR319, Pgi-miR172, and Pgi-miR164 also changed to varying degrees, suggesting that MeJA induction is associated with changes in particle levels, active components, and miRNA expression characteristics of ginseng-derived exosome-like nanovesicles.

In summary, this study preliminarily established a method for the extraction and characterization of ginseng-derived exosome-like nanovesicles and revealed the changing characteristics of vesicle-related particles and molecular composition under MeJA induction conditions, providing experimental evidence and methodological reference for further scalable preparation, component regulation, and biological function research of ginseng-derived exosome-like nanovesicles.

## Author Contributions

Leyi XIAO: Conceptualization; methodology. Yuanzhu LIU: funding acquisition. Chi XIONG: writing—original draft; software; for malanalysis. Cheng SUN: supervision. Junquan CHENG: Visualization; investigation. Junxia XIAO: resources. Yuping RONG: resources. Rongchun HAN: resources. Qinying YANG: resources. Xiao LIU: resources; project administration. Rui LI: resources. Cuiling ZHANG: Visualization; resources. Xin WAN: Visualization; investigation. Xianqian LIU: Visualization; investigation. Jihong HUANG: writing— review and editing; resources; funding acquisition. All authors have read and agreed to the published version of the manuscript.

## Data Availability

Data will be made available on request.

## Acknowledgments

We gratefully acknowledge all contributors to this work.

## Supporting Information

The following is the Supplementary data to this article:

Supplementary Table S2: Analysis of Extracellular vesicle like Liposomes Derived from Culture Medium.

Supplementary Table S3: Lipid analysis of extracellular vesicle like nano vesicles derived from tissue cultured adventitious roots.

